# Brodifacoum does not modulate human cannabinoid receptor-mediated hyperpolarization of AtT20 cells or inhibition of adenylyl cyclase in HEK 293 cells

**DOI:** 10.1101/589341

**Authors:** Shivani Sachdev, Rochelle Boyd, Natasha L Grimsey, Mark Connor

## Abstract

**Background:** Synthetic cannabinoids are a commonly used class of recreational drugs that can have significant adverse effects. There have been sporadic reports of co-consumption of illicit drugs with rodenticides such as warfarin and brodifacoum (BFC) over the past 20 years but recently, hundreds of people have been reported to have been poisoned with a mixture of synthetic cannabinoids and BFC. We have sought to establish whether BFC directly affects cannabinoid receptors, or their activation by the synthetic cannabinoid CP55940 or the phytocannabinoid Δ^9^-tetrahydrocannabinol (Δ^9^-THC).

**Methods:** The effects of BFC on the hyperpolarization of wild type AtT20 cells, or AtT20 cells stably expressing human CB_1_- and CB_2_-mediated receptors, were studied using a fluorescent assay of membrane potential. The effects of BFC on CB_1_ and CB_2_ mediated inhibition of forskolin-stimulated adenylyl cyclase (AC) activation was measured using a BRET assay of cAMP levels in HEK 293 cells stably expressing human CB_1_ and CB_2_.

**Results:** BFC did not activate CB_1_ or CB_2_ receptors, or affect the hyperpolarization of wild type AtT20 cells produced by somatostatin. BFC (10 *µ*M) did not affect the hyperpolarization of AtT20-CB_1_ or AtT20-CB_2_ cells produced by CP55940 or Δ^9^-THC. BFC (1 *µ*M) did not affect the inhibition of forskolin-stimulated AC activity by CP55940 in HEK 293 cells expressing CB_1_ or CB_2_. BFC (1 *µ*M) also failed to affect the desensitization of CB_1_ and CB_2_ signalling produced by prolonged (30 min) application of CP55940 or Δ^9^-THC to AtT20 cells.

**Discussion:** BFC is not a cannabinoid receptor agonist, and appeared not to affect cannabinoid receptor activation. Our data suggests there is no pharmacodynamic rationale for mixing BFC with synthetic cannabinoids, however, it does not speak to whether BFC may affect synthetic cannabinoid metabolism or biodistribution. The reasons underlying the mixing of BFC with synthetic cannabinoids are unknown, and it remains to be established whether the “contamination” was deliberate or accidental. However, the consequences for people who ingested the mixture were often serious, and sometimes fatal, but this seems unlikely to be due to BFC action at cannabinoid receptors.

## Introduction

Brodifacoum (BFC) is an inhibitor of vitamin K epoxide reductase and active ingredient of rodenticides (Brekenridge et al., 1978). There have been sporadic reports of brodifacoum consumption with drugs such as cocaine and cannabis (La Rosa et al., 1997; Waien et al, 2001; Spahr et al., 2007), and recently a large number of people have been hospitalized with poisoning by brodifacoum and related compounds following ingestion of what are believed to be synthetic cannabinoid receptor agonists (SCRA; Kelkar et al., 2018; Riley et al., Mortiz et al., 2018, Panigrahi et al., 2018). There is a little evidence suggesting that occasionally people have deliberately combined brodifacoum with cannabis (La Rosa *et al.*, 1997; Spahr *et al.*, 2007), and the apparent mixing of brodifacoum with a variety of different SCRA could suggest a deliberate attempt to enhance the effects of the drugs by recruiting either a pharmacokinetic or pharmacodynamic mechanism. In this study, we have examined the effects of brodifacoum on the acute signalling of human CB_1_ and CB_2_ receptors in AtT20FlpIn and HEK 293 cells. In AtT20FlpIn cells, activation of heterologously expressed CB_1_ or CB_2_ produces a hyperpolarization, mediated by activation of G protein-gated inwardly rectifying K channels (Mackie et al., 1995; Banister *et al.*, 2016). In HEK 293 cells, we measured the real time modulation of forskolin-stimulated cAMP accumulation (Cawston *et al.*, 2013). We found that cannabinoid-induced signaling was not affected by brodifacoum, indicating that combining SCRA with brodifacoum is not likely to enhance user experience through interactions with cannabinoid receptors.

## Methods

### Hyperpolarization Assay

Experiments on AtT20FlpIn cells stably transfected with human CB_1_ (AtT20-CB_1_) or CB_2_ (AtT20-CB_2_) were carried out essentially as described in Banister *et al.*, 2016. The AtT20FlpIn cells were created in our laboratory from wild type AtT20 cells we purchased from the American Type Culture Collection (ATCC CRL-1795). The assay method is based on that outlined in detail in Knapman *et al.*, 2013. Cells were grown in DMEM (#D6429, Sigma-Aldrich, Castle Hill, NSW) supplemented with 10% fetal bovine serum (FBS, #12003C, SAFC Biosciences, Brooklyn, Vic), 100 units penicillin/100 *µ*g ml^−1^ streptomycin (1%, #15140122 Life Technologies, Scoresby, Vic), hygromycin gold (80 *µ*g ml^−1^, #ant-hg, Invivogen, San Diego, CA). Cells were grown in 75 cm^2^ flasks and passaged when 80-90% confluent. On the evening before experiments, cells were detached using trypsin/EDTA solution (#T3924, Sigma-Aldrich), resuspended in L-15 media (#11415064, Life Technologies) supplemented with 1% FBS, penicillin/streptomycin, and glucose (15 mM, SIGMA #G7021) and plated onto 96 well black walled, clear bottomed, culture plates which had been previously coated with poly-D-lysine (SIGMA #P6407). Cells were incubated overnight in a humidified incubator in room air.

Proprietary FLIPR membrane potential dye (blue, #R8034, Molecular Devices, Sunnyvale CA) was dissolved in HEPES-buffered saline (HBS) of composition (mM) NaCl 145, HEPES 22, Na_2_HPO_4_ 0.338, NaHCO_3_ 4.17, KH_2_PO_4_ 0.441, MgSO_4_ 0.407, MgCl_2_ 0.493, CaCl_2_ 1.26, glucose 5.56 (pH 7.4, osmolarity 315 ± 15) and added to the cells an hour before fluorescence reading began. Dye was used at 50% of the manufacturers recommended concentration, and cells were incubated at 37 °C in humidified room air for loading. Plates were read using a Flexstation 3 (Molecular Devices) plate reader at 37 °C. Plates were excited at a wavelength of 530 nm, emission was measured at 565 nm, with cut-off filter at 550 nm. Drugs were added using the pipetting function of the Flexstation in a volume of 20 *µ*l after recording 60-120 s of baseline fluorescence. Readings were made every 2 seconds. Drug stocks were made up in DMSO (#D8418, Sigma-Aldrich) and diluted on the day of experiment, the final concentration of DMSO in the assay was 0.1 %.

Data were expressed as the percentage change in baseline fluorescence produced by drug addition. The change in fluorescence produced by vehicle addition was subtracted from the traces before this calculation. Data is expressed as the mean ± SEM of at least 5 independent determinations performed in duplicate, unless otherwise noted. Pooled data was fit to a 4 parameter logistic equation in Graphpad PRISM 7 (GraphPad Software, San Diego CA).

### Assay of cAMP levels

Human embryonic kidney (HEK) 293 FlpIn cells stably transfected with human CB_1_ or CB_2_ receptors tagged with three haemagglutinin epitopes at the amino terminus and human G protein gated inwardly rectifying potassium channel 4 (GIRK4) were used (the construction of these cells will be described in another place, and we did not assay CB receptor coupling to GIRK4 in this study). Cells were grown in DMEM containing 10% FBS and 100 units/penicillin, 100 μg/ml streptomycin and were maintained under selection with hygromycin (80 *µ*g ml^−1^) and G418 (400*µ*g ml^−1^). HEK 293 FlpIn cells were originally obtained from Life Technologies (now Thermofisher, #75007).

Cellular cAMP levels were measured using the pcDNA3L-His-CAMYEL plasmid, which encodes the cAMP sensor YFP-Epac-RLuc (CAMYEL), (Cawston *et al.*, 2013; Hunter *et al.*, 2017). The pcDNA3L-His-CAMYEL was a kind gift from Dr. Angela Finch (The University of New South Wales, NSW, Australia), and originally obtained from American Type Culture Collection (Manassas, VI, USA). Cells were seeded in 10 cm dishes at a density of 6,000,0000 such that they would be 60-70% confluent the next day. The day after seeding, pcDNA3L-His-CAMYEL plasmid were transiently transfected into cells using linear polyethyleneimine (PEI, m.w. 25 kDa) (#23966, Polysciences, Warrington, PA, USA). The DNA-PEI complex mixture was added into the cells at the ratio of 1:6, and incubated for 24 hours in 5% CO_2_ at 37 °C. After the incubation, cells were detached from the dish using trypsin/EDTA and the pellet was resuspended in 10 ml Leibovitz’s L-15, no phenol red (#21083027, Gibco) media supplemented with 1% FBS, 100 units/penicillin, 100 μg/ml streptomycin and 15 mM glucose. The cells were seeded at a density of 100,000 cells per well in poly D-lysine (Sigma-Aldrich) coated, white wall, clear bottom 96 well microplates. Cells were incubated overnight at 37 °C in ambient CO_2_.

On the following day, drugs were prepared in HBS containing 0.1 mg ml^−1^ BSA. For measurement of cAMP inhibition, all the drugs were made in 3 *µ*M of forskolin. Coelenterazine-h substrate (2.5 *µ*M) (#S2011, Promega, Madison, WI, USA) was added to the cells, and incubated for 5 mins prior to the addition of drugs or vehicle. Luminescence was measured using a Flexstation 3 (Molecular Devices) microplate reader at 37 °C. The cells were measured at an emission wavelength of 461 nm and 542 nm simultaneously, with an integration time of 1 s. Drugs were added in a volume of 10 *µ*l (10X) to each well to give the desired concentration. The final concentration of DMSO in each well was always 0.1%. Raw data are presented as inverse BRET ratio of emission at 461/542. Background reading (no substrate) was subtracted from raw values before calculating ratios. For convenience, values are expressed such that an increase in ratio correlates with increase in cAMP production. AUC analysis was performed in GraphPad prism (Graph Pad Software Inc., San Diego, CA), and data were expressed as percentage of the difference between basal (vehicle, 0%) and forskolin (100%) values over a 5 minute period after forskolin addition.

(-) CP 55940 was from Cayman Chemical (#90084, Ann Arbor MI), Δ^9^-tetrahydrocannabinol (THC) was from THCPharm (Frankfurt, Germany) and was a kind gift from the Lambert Initiative for Cannabis Therapeutics (University of Sydney). Brodifacoum was from Sigma-Aldrich (#46036), and forskolin was from Ascent Scientific Ltd.

Data was normally distributed (D’Agonstino and Pearson normality test, PRISM), differences between groups were tested using unpaired Student’s t-Test (PRISM).

## Results

Application of brodifacoum for 5 minutes at concentrations up to 30 *µ*M did not significantly affect the fluorescence of AtT20 cells expressing CB_1_ or CB_2_ receptors (Figure 1). Prolonged exposure to brodifacoum at concentrations greater than 10 *µ*M produced decreases in fluorescence in AtT20 cells expressing CB receptors as well as wild type cells, and so for experiments examining the potential interaction between brodifacoum and cannabinoids we used a concentration of 1 *µ*M.

**Figure 1:**
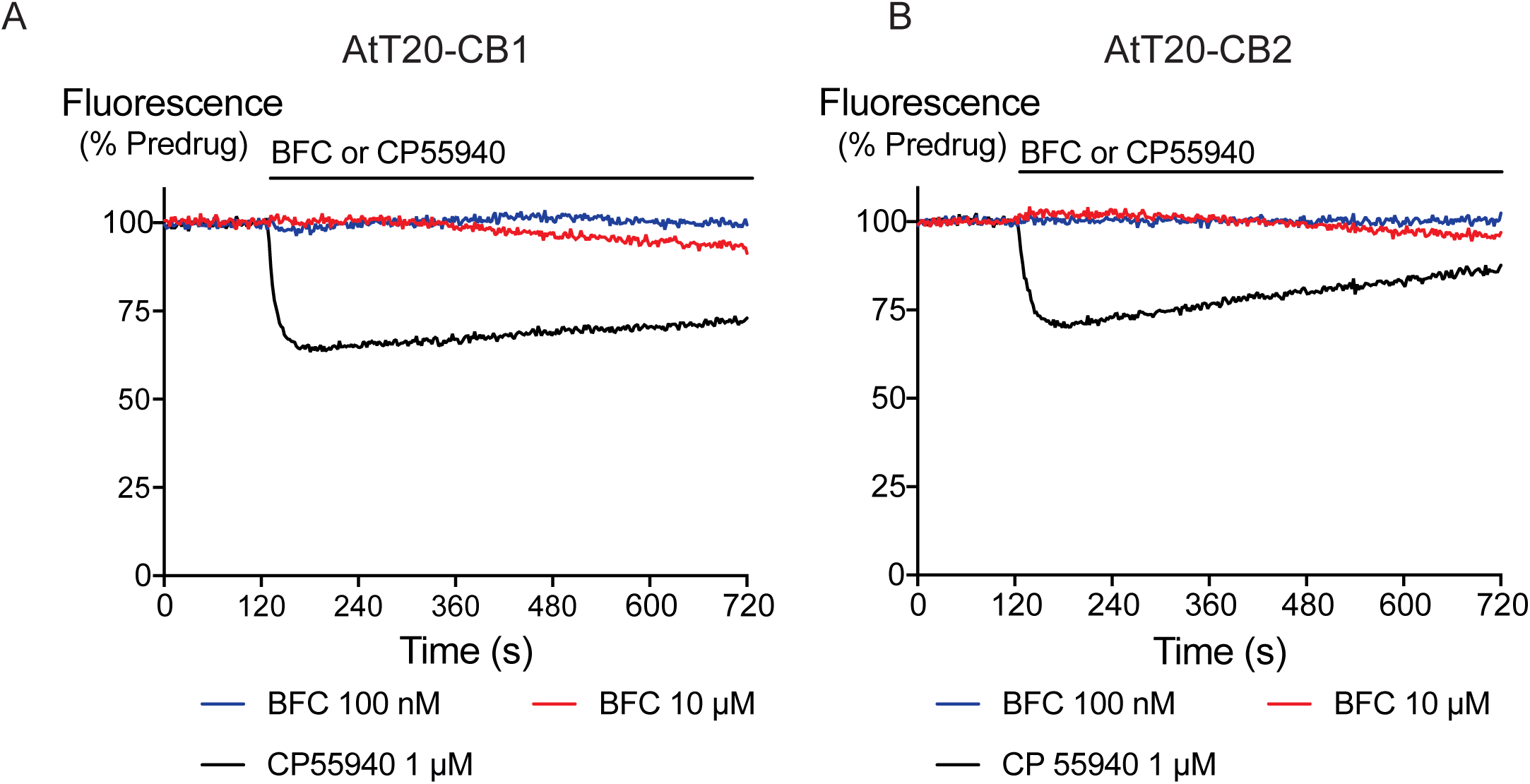
The effects of brodifacoum (BFC) and CP55940 in AtT20 cell expressing CB_1_ or CB_2_. Representative traces showing the change in fluorescence induced by application of CP55940 (1 *µ*M) but not BFC (10 *µ*M) in **A)** AtT20-CB_1_ and **B)** AtT20-CB_2_ cells. Values are expressed as a percentage of predrug baseline. A reduction in fluorescence indicates a hyperpolarization. The prolonged application of BFC (10 *µ*M) produces small changes in the fluorescence in AtT20 cells expressing cannabinoid receptors. Drug was added for the duration of the bar; the traces are representative of at least five independent experiments.

We generated concentration-response curves for the high efficacy cannabinoid agonist CP55940 and the lower efficacy agonist THC after 5 minutes of exposure to brodifacoum (Figure 2). In AtT20-CB_1_ cells, application of CP55940 produced a maximum change in fluorescence of 33 ± 1 %, with a *p*EC_50_ of 7.7 ± 0.04; in the presence of brodifacoum the maximum change in fluorescence of 33 ± 1 %, with a pEC50 of 7.7 ± 0.06. In AtT20-CB_2_ cells, application of CP55940 produced a maximum change in fluorescence of 29 ± 1.1 %, with a *p*EC_50_ of 7.3 ± 0.1; in the presence of brodifacoum the maximum change in fluorescence of 31 ± 1.2 %, with a *p*EC_50_ of 7.4 ± 0.1 (Figure 2). Brodifacoum failed to affect the hyperpolarization produced by THC in AtT20-CB_1_ cells (control, *p*EC50 6.4 ± 0.6, maximum change in fluorescence 18 ± 5 %; in brodifacoum, *p*EC_50_ 6.5 ± 0.5, max 18 ± 5 %). In AtT20-CB_2_ cells THC produces a small hyperpolarization, the response to 10 *µ*M THC was unchanged in the presence of brodifacoum (6.4 ± 1.2 % in control, 7.4 ± 1.8 % in brodifacoum, P = 0.65) (Figure 2). Application of brodifacoum (10 *µ*M) or CP55940 (1 *µ*M) for 5 minutes produced very small changes in the fluorescence of wild type AtT20 cells, and neither drug affected the response to subsequently applied somatostatin (100 nM), which activates native sst receptors in AtT20 cells (Gunther *et al.*, 2016) (Supplementary Figure 1).

**Figure 2:**
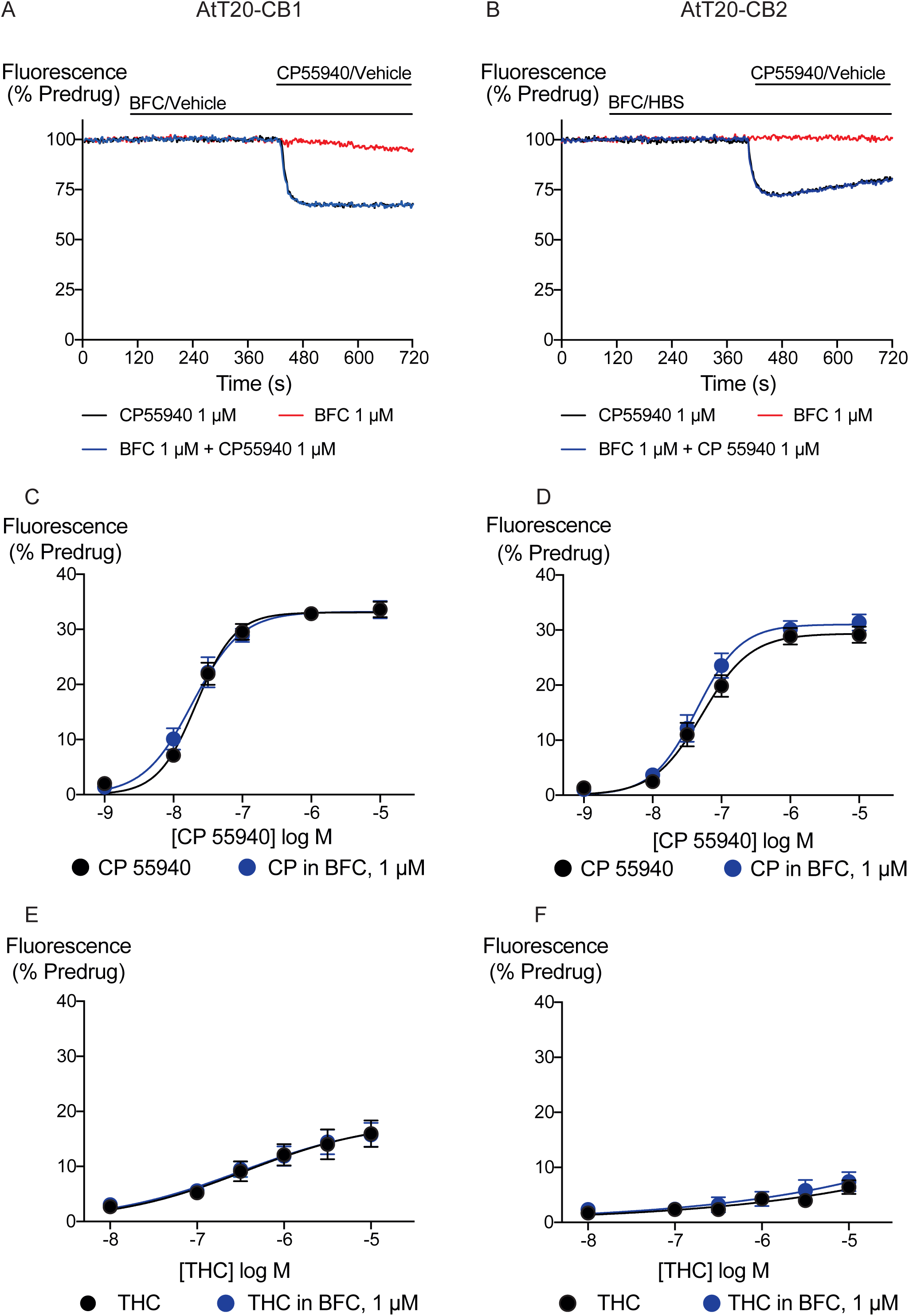
Brodifacoum (BFC) effect on CP55940 and Δ^9^-THC induced hyperpolarization of AtT20 cell expressing CB_1_ or CB_2_. Representative traces showing the change in fluorescence for CP55940 on **A)** AtT20-CB_1_, and **B)** AtT20-CB_2_ in the presence of BFC 1 *µ*M or vehicle. Values are expressed as a percentage of predrug baseline. A reduction in fluorescence indicates a hyperpolarization. Drugs were added for the duration of the bar; the traces are representative of at least five independent experiments. Concentration response curve of hyperpolarization of AtT20-CB_1_ or AtT20-CB_2_ cells stimulated with **C)**, **D)** CP55940 or **E)**, **F)** Δ^9^-THC in the continued presence of either HBS or BFC. Data represents the mean ± SEM of five independent determinants performed in duplicate. There was no difference in the potency or maximal effect of CP55940 and Δ^9^-THC between HBS or in presence of BFC.

Inhibition of adenylyl cyclase activity is another significant biological effect of cannabinoid receptor activation. Brodifacoum (300 nM – 30 *µ*M) co-applied with forskolin (3 *µ*M) for 10 minutes did not affect increases in cAMP levels in HEK 293 cells expressing CB_1_ or CB_2_ (Figure 3). Brodifacoum (1 *µ*M) incubation for 5 minutes also failed to affect the CP55940 inhibition of forskolin-stimulated cAMP elevation. In cells expressing CB_1_, CP55940 inhibited cAMP with a *p*EC_50_ of 7.5 ± 0.3, to a minimum of 52 ± 12 % of forskolin alone; in the presence of brodifacoum these were *p*EC_50_ 7.4 ± 0.2 and minimum of 52 ± 7% of the forskolin response. Brodifacoum also did not affect forskolin-stimulated cAMP levels HEK293 cells expressing CB_2_ (Figure 3), or CP55940 inhibition of cAMP levels (*p*EC_50_ in control cells expressing CB_2_ 7.4 ± 0.2, to a minimum of 39 ± 7 %; in brodifacoum *p*EC_50_ of 7.5 ± 0.1; to a minimum of 45 ± 4 %).

**Figure 3:**
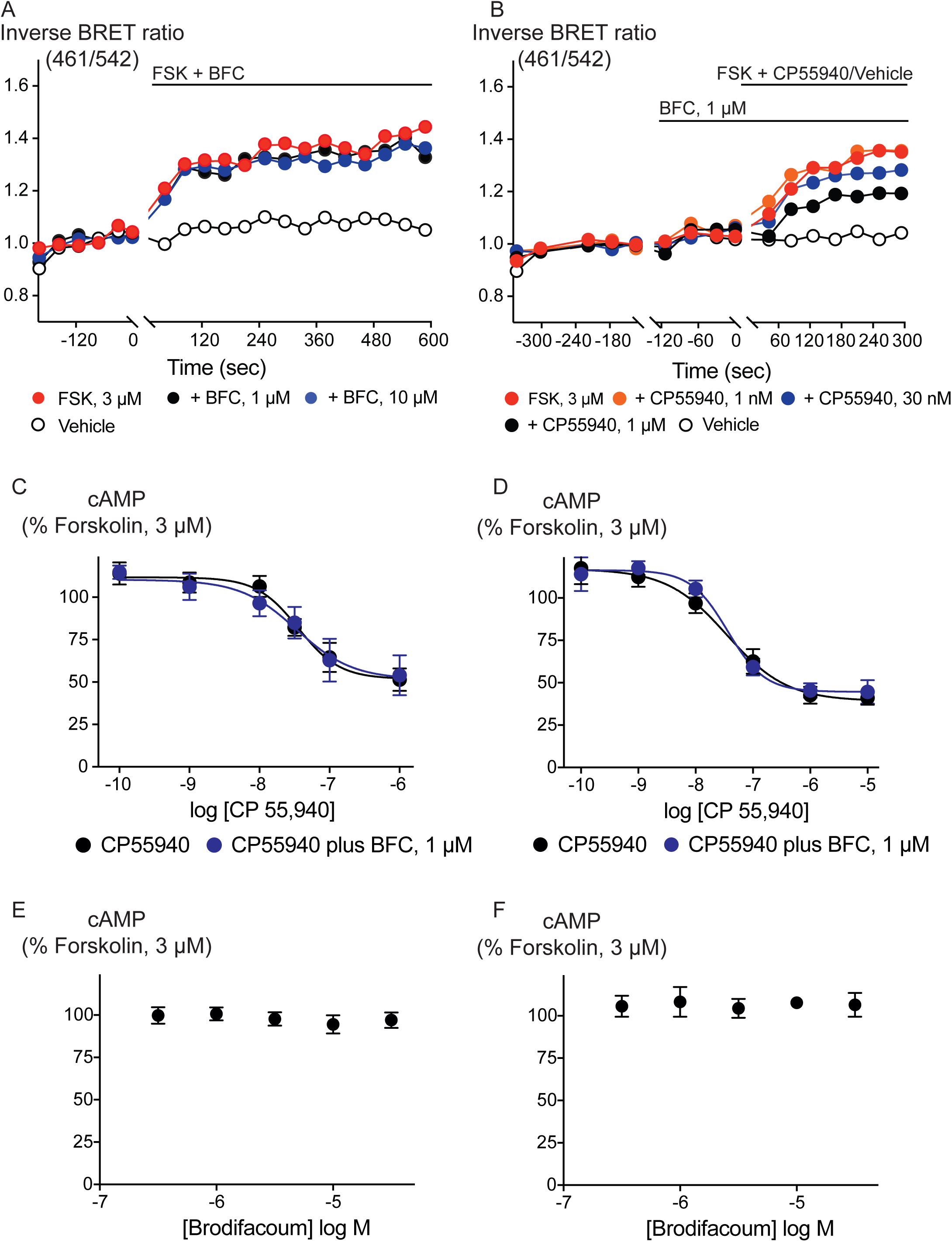
Brodifacoum (BFC) does not modulate cAMP accumulation via CB_1_ or CB_2_ receptors expressed in HEK 293 cells. Representative data from the CAMYEL assay for HEK 293 cells expressing CB1 receptors, an increase in inverse BRET ratio (emission at 461/542 nm) corresponds to an increase in cAMP. **A)** BFC does not affect the rapid increase in cAMP production produced by forskolin (3 *µ*M); **B)** BFC (1 *µ*M) does not affect responses to forskolin (3 *µ*M) applied in the presence of CP55940. Data are representative of at least five independent experiments. Concentration response curve showing CP55940 induced inhibition of forskolin-stimulated cAMP elevation in presence and absence of BFC 1 *µ*M on HEK 293 cells expressing **C)** CB_1_ or **D)** CB_2_. Data are expressed as a percentage of response produced by forskolin (3 *µ*M), and plotted as mean ± SEM of five independent determinants performed in duplicate. Concentration response curve showing the effect of BFC on forskolin (3 *µ*M)-stimulated cAMP elevations in HEK 293 cells expressing **E)** CB_1_ or **F)** CB_2_. Data are expressed as a percentage of response produced by forskolin (3 *µ*M), and plotted as mean ± SEM of five independent determinants performed in duplicate.

We also examined the possibility that brodifacoum could affect the sustained responses to CP55940 or THC. As previously described (Cawston *et al.*, 2013), prolonged application of cannabinoids in AtT20-CB_1_ cells produces a response that wanes over time, reflecting desensitization of receptor signaling. The degree to which this desensitization reflects changes in signaling specific to cannabinoid receptors is tested by application of somatostatin, which activates receptors native to AtT20 cells (Gunther *et al.*, 2016; Heblinski *et al.*, 2019). In these experiments, CP55940 (100 nM) or THC (10 *µ*M) were applied 2 minutes after addition of brodifacoum (1 *µ*M), and the fluorescence monitored for 30 minutes before the addition of somatostatin (100 nM) (Figure 4). Desensitization was quantified after 30 min of agonist application, and was expressed as the % decline from the peak response. We did not observe any significant difference in the desensitization of CB_1_ signaling mediated by CP55940 (100 nM) when co-applied with brodifacoum (P = 0.55). The presence of brodifacoum had no effect on the SOMATOSTATIN (100 nM) induced hyperpolarization alone, or after 30 mins of CP55940 treatment (P = 0.75)(Supplementary Figure 2). The desensitization produced by THC (10 *µ*M, 30 mins) in AtT20-CB_1_ cells was not different when co-applied with brodifacoum, (Control, 65 ± 6%; brodifacoum treated, 53 ± 8%, P = 0.3) (Figure 4). A similar reversal of the hyperpolarization produced by CP55940 (100 nM) in AtT20-CB_2_ cells was also observed. Treatment with brodifacoum did not significantly affect the desensitization produced by CP55940 compared to control cells (Control, 77 ± 6%; brodifacoum treated, 63 ± 8%, P = 0.2). THC (10 *µ*M, 30 mins) signaling at CB_2_, although modest, also declined during continuous drug exposure, and this was also not affected by co-application of brodifacoum (37 ± 14% in control, 20 ± 7% in brodifacoum treated, P = 0.3)(Figure 4). The hyperpolarization induced by SOMATOSTATIN after prolonged application of CP55940 (P=0.56) or THC (P=0.87) to AtT20-CB_2_ cells was also not significantly different in the presence of brodifacoum (Supplementary Figure 2).

**Figure 4:**
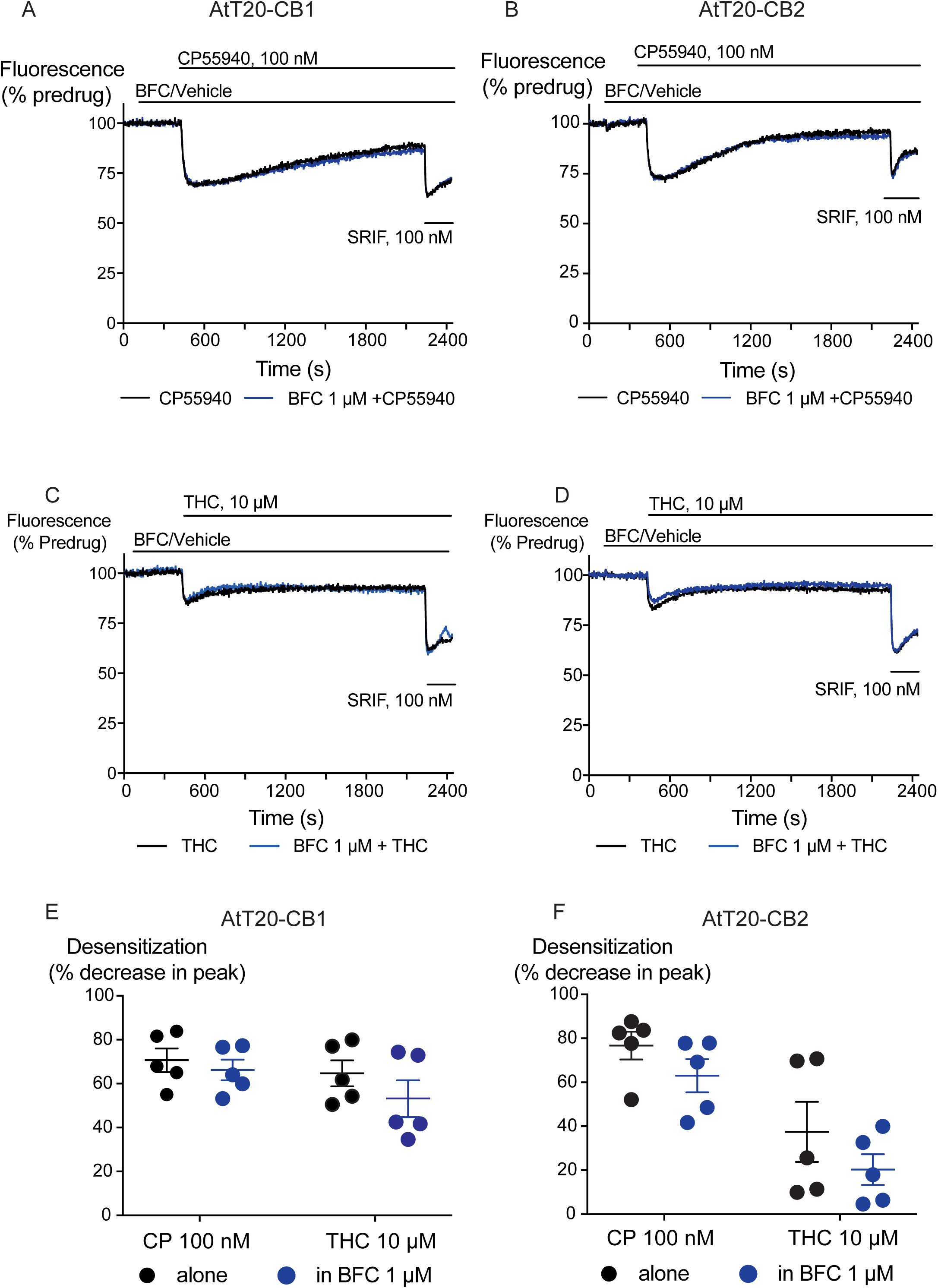
The effect of brodifacoum (BFC) on CP55940 and Δ^9^-THC mediated desensitization of signalling in AtT20-CB_1_ and -CB_2_. Representative traces showing desensitization of signalling in AtT20-CB_1_ and AtT20-CB_2_ on prolonged stimulation with **A)**, **C)** CP55940 (100 nM) or **B)**, **D)** Δ^9^-THC (10 *µ*M) in the presence of BFC 1 *µ*M or HBS. Cells were challenged with somatostatin (100 nM) after 30 minutes of CP55940 or Δ^9^-THC. Drugs were added for the duration of the bar; the traces are representative of at least five independent experiments. Scatter dot plot representing desensitization of **E)** CB_1_ and **F)** CB_2_ on exposure to CP55940 or Δ^9^-THC for 30 mins in the presence of BFC 1 *µ*M or HBS. This plot shows percentage desensitization comparing peak fluorescence after the addition of drugs and 30 mins post addition. Data represents the mean ± SEM of five independent determinants performed in duplicate.

## Discussion

The principal finding of this study is that brodifacoum does not affect CB_1_ or CB_2_ signalling, either to K channels in AtT20 cells or adenylyl cyclase in HEK 293 cells. In the assay of K channel activation, there was no effect on the concentration response relationship for CP55940 or THC, and brodifacoum did not affect the desensitization of signalling produced by prolonged application of either drug. Activation of GIRK is mediated by the Gβγ subunits of G protein heterotrimers, and many Gi/Go coupled receptors effectively signal through this pathway in AtT20 cells (e.g. Mackie et al., 1995; Gunther et al., 2016, Knapman et al., 2013, Heblinski et al., 2018). We have previously used the fluorescent measurement of membrane potential to study CB_1_ and CB_2_ agonists, antagonists, and allosteric modulators of CB_1_ (Cawston *et al.* 2013). Brodifacoum had no effect on the potency, maximum effect or time-dependence of the actions of the high efficacy synthetic cannabinoid CP55940 or the lower efficacy phytocannabinoid THC, indicating that it is unlikely to act as modulator of the pharmacodynamic effects of cannabinoids.

Inhibition of adenylyl cyclase activity by CB receptors is mediated via the Gα subunits of G protein heterotrimers, and brodifacoum also failed to affect this signal transduction pathway. The precise cellular signalling mechanisms responsible for the subjective effects of *Cannabis* and synthetic cannabinoid agonists are not established, although the signal transduction of cannabinoid receptors has been extensively studied (Howlett & Abood, 2017; Ibsen et al. 2017) and it is unlikely that any one pathway is responsible. It remains formally possible that brodifacoum could selectively modulate pathways other than Gβγ-mediated activation of GIRK or Gα-mediated inhibition of cAMP accumulation, but the lack of any effect whatsoever on the effects of CP55940 or THC suggest that ligand interactions with cannabinoid receptors are unaffected by brodifacoum.

The concentration of brodifacoum in blood or brain after co-ingestion with synthetic cannabinoids is unknown. However, concentrations of up to 3 *µ*M have been reported in the serum of people who have deliberately ingested large quantities of rat poison (Weitzel et al., 1990; Hollinger et al., 1993). Brodifacoum at 1 *µ*M failed to affect CB_1_ or CB_2_ receptor signalling when measured continuously over a period of 30 minutes, and 10 *µ*M brodifacoum failed to mimic or affect the acute response to a maximally effective concentration CP 55940, although at this concentration prolonged application of brodifacoum produced a decrease in the fluorescence of wild type AtT20 cells, as well as those expressing CB_1_ and CB_2_ receptors. This effect at higher concentrations may reflect direct interactions of brodifacoum with cell membranes (Maragoni *et al.*, 2016). Concentrations of brodifacoum in the upper range of what we tested are achieved only after ingestion of large amounts of rat bait, it is possible that they could be achieved while ingesting contaminated synthetic cannabinoids, but this remains unreported.

Several case reports suggest an interaction between therapeutic warfarin and cannabis or cannabidiol (Grayson *et al.*, 2018; Yamreudeewong *et al*., 2009). It has been suggested that cannabinoid inhibition of enzymes responsible for the metabolism of warfarin can increase blood levels of the drug, and while these studies have focussed on potentially dangerous changes in warfarin concentration, levels of cannabinoids could also be reciprocally elevated. Such interactions may inform a decision to deliberately combine “superwarfarin” with SCRA, as has been previously suggested for cannabis (La Rosa *et al* 1997; Spahr *et al.*, 2007), although whether brodifacoum is metabolized by pathways shared with SCRA in humans is unknown. Apart from the obvious danger of ingesting brodifacoum, altering the metabolism of SCRA is likely to have unpredictable consequences, as some metabolites of SCRA retain cannabinoid receptor activity (e.g. Brents *et al.*, 2011; Chimalaconda *et al.*, 2012; Longworth *et al.*, 2017) and may contribute to the overall SCRA experience.

Ingestion of brodifacoum is relatively common, while death from exposure is rare, owing to ready treatment with vitamin K (King and Tran, 2015; Gummin *et al.*, 2018). The high number of deaths associated with the combination of SCRA and anticoagulants in 2018 (at least 8, Connors, 2018) may point to an interaction between the drugs or a very high dose of ingested anticoagulant. It may also reflect the identity and dose of the synthetic cannabinoid(s) consumed (which remains largely unreported), as well as the general health status of the drug users. Deaths from synthetic cannabinoid exposure are uncommon, but well documented (e.g. Kasper et al., 2018; Treki et al., 2015). While there is a general acceptance that brodifacoum or a similar agent is responsible for the coagulopathies associated with synthetic cannabinoid ingestion, identification of the synthetic cannabinoid has not been reported in most cases. Intriguingly, several groups have reported cannabinoid receptor ligands based on a coumarin scaffold (Behrenswerth *et al.*, 2009; Han *et al.*, 2015). While these drugs have been reported to be either antagonists/inverse agonists (Behrenswerth *et al.*, 2009) or CB_2_-selctive agonists (Han et al., 2015), they remain largely uncharacterized. Given the propensity of chemists producing and in some cases designing cannabinoids for the recreational market, it cannot be ruled out that some of the coagulopathy associated with synthetic cannabinoid use arises from a novel, coumarin-based cannabinoid that retains some of the vitamin K epoxide inhibitory of warfarin and brodifacoum.

In conclusion, we report that brodifacoum does not appear to be an agonist or antagonist of human cannabinoid receptors, and it also does not appear to be an allosteric modulator of CB_1_ or CB_2_ activation of K channels or inhibition of adenylyl cyclase. Why brodifacoum has been mixed with synthetic cannabinoid receptor agonists remains a matter for speculation, although an intended effect on synthetic cannabinoid drug pharmacokinetics cannot be ruled out.

## Acknowledgements

This work was supported by the NHRMC of Australia (APP1107088). SS was supported by an International Research Excellence Scholarship from Macquarie University. We thank Dr Sam Banister for his insight, especially around the possibility of poisoning with coumarin-derived synthetic cannabinoids, and Dr Marina Santiago, for her advice on the cAMP assay. RB (GIRK), SS (GIRK and cAMP) performed experiments and analysed data. NG created the HEK293 cells and provided invaluable input into the design and analyses of the cAMP experiments. MC conceived the paper, analysed data and wrote the Ms with RB and SS, all authors have seen and approved the final version. The authors declare they have no conflicts of interest with regard to this work.

## Supplementary Figures

**Supplementary Figure 1:**
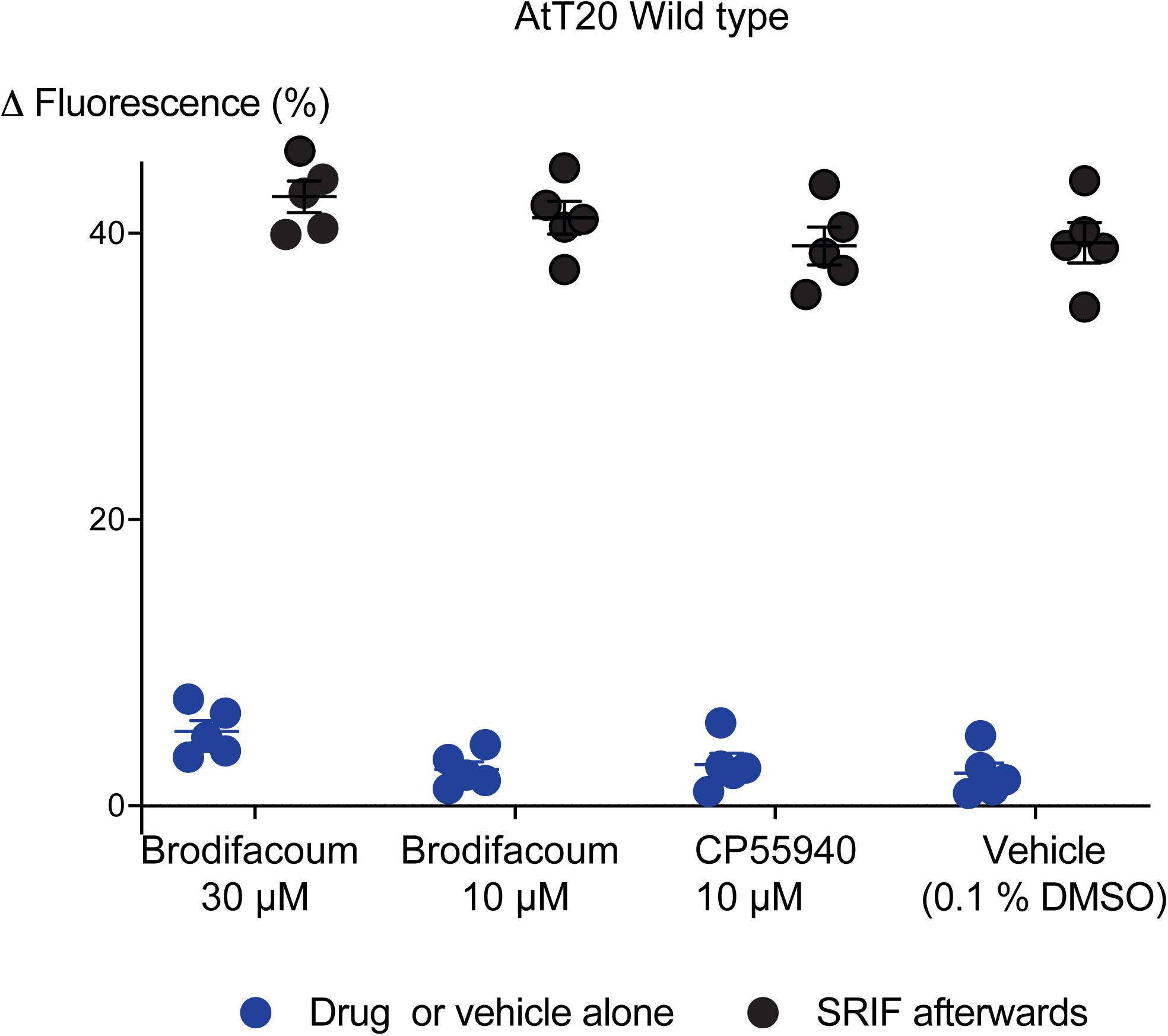
Effects of brodifacoum (BFC) and CP55940 in wild type AtT20 cells. Scatter dot plot representing the percentage change in fluorescence for BFC (30 *µ*M), BFC (10 *µ*M), CP55940 (10 *µ*M), and Vehicle (0.1% DMSO) alone (blue dots), and the response to the subsequent addition of SOMATOSTATIN (100 nM) to AtT20-WT cells (black dots). Data represents the mean ± SEM of five independent determinants performed in duplicate (p > 0.05).

**Supplementary Figure 2:**
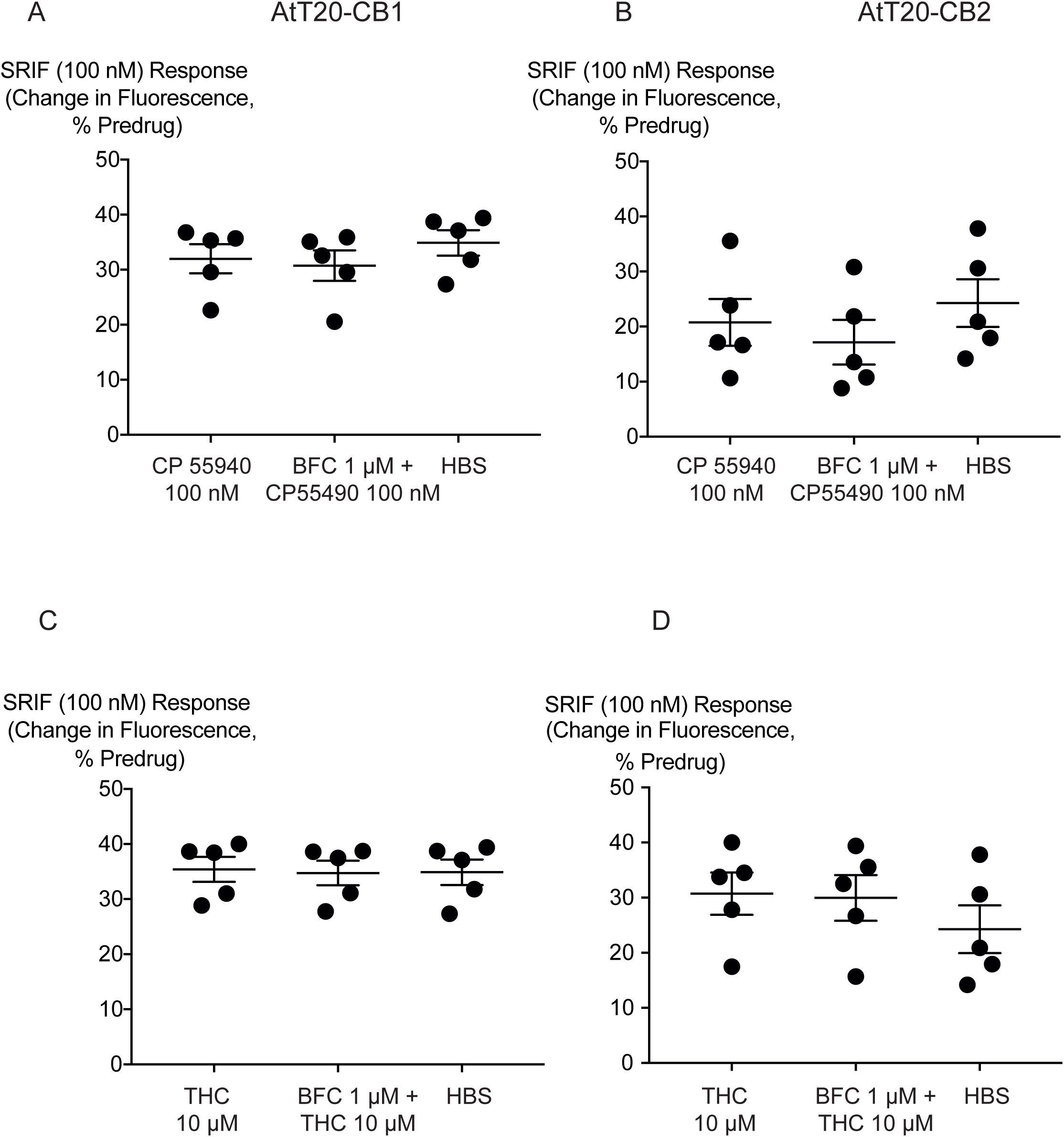
Effect of brodifacoum (BFC) on somatostatin (SRIF) challenge after 30 minutes of drugs on AtT20-CB_1_ and -CB_2_ cells. Comparison of percentage change in fluorescence after SRIF (100 nM) challenge on AtT20-CB_1_, and AtT20-CB_2_ in the continuous presence of **A)**, **C)** CP55940 or **B)**, **D)** Δ^9^-THC added with either HBS or BFC (1 *µ*M). BFC did not affect the hyperpolarization induced by SRIF after prolonged application of CP55940 or Δ^9^-THC. Data represents the mean ± SEM of five independent determinants performed in duplicate (p > 0.05).

